# Intrinsic noise modulation in closed oligomerization-type systems^⋆^

**DOI:** 10.1101/254763

**Authors:** Marianne Rooman, Fabrizio Pucci

## Abstract

How random fluctuations impact on biological systems and what is their relationship with complexity and energetic cooperativity are challenging questions that are far from being elucidated. Using the stochastic differential equation formalism, we studied analytically the effect of fluctuations on a series of oligomerization processes, in which several molecules of the same or different species interact to form complexes, without interaction with the environment. The conservation of the total number of molecules within the systems imposes constraints on the stochastic quantities, among which the negativity of the covariances and the vanishing of the determinant of the covariance matrix. The intrinsic noise on the number of molecules of each species is represented by the Fano factor, defined as the variance to mean ratio. At the equilibrium steady states, the sum of the Fano factors of all molecular species is equal to the rank of the system, independently of the parameters. The Fano factors of the individual molecular species are, however, parameter dependent. We found that when the free energy cooperativity of the reactions increases, the intrinsic noise on the oligomeric product decreases, and is compensated by a higher noise on the monomeric reactants and/or intermediate states. The noise reduction is moreover more pronounced for higher complexity systems, involving oligomers of higher degrees.

## 1. INTRODUCTION

Although stochastic fluctuations play a major role in the dynamics of many biological processes, they are often overlooked. Besides the fundamental challenge of understanding how biosystems deal with these fluctuations, the ability to control them would have valuable applications in synthetic biology in view of rationally designing systems with a precise number of molecules of each species or, on the contrary, with very noisy behaviors.

To obtain a proper description of these effects, stochastic modeling approaches are required (1; 2; 3). However, the large size and complexity of biological systems make them difficult to model mathematically, especially via stochastic simulations in which the parameter space becomes rapidly too large to be tractable. This has up to now prevented getting general insights into the biological impact of stochastic fluctuations.

An intriguing open question concerns the relationship between the complexity of biological systems and the modulation of the intrinsic noise (4; 5; 6). In a recent study (7), we have made an important step towards this goal. We analyzed a large series of chemical reaction networks with different topologies and degrees of complexity, interacting or not with the environment, and showed that the global level of intrinsic noise at the steady state is directly related to the network’s structure. For networks with a detailed-balanced (equilibrium) or complex-balanced steady state, the global noise level is constant, whereas for other homo-oligomerization networks, the global fluctuation level decreases or increases according to whether the fluxes between molecular species flow from low to high, or from high to low, complexity.

In this paper, we studied in detail a subset of these networks and more specifically, closed biological systems representing molecular homo-or hetero-oligomerization processes, which admit an equilibrium steady state. Although the global intrinsic noise is in this case constant for all parameter values, we see that noise reduction or amplification occurs for some of the molecular species in a parameter-dependent fashion.

## 2. MATHEMATICAL MODELING USING ITŌ STOCHASTIC DIFFERENTIAL EQUATIONS

Let us consider closed systems consisting of several molecular species that interact with each other to form molecular complexes. These molecules are, for example, proteins that oligomerize or that bind to ligands, DNA or RNA. As biological systems are inherently stochastic, their variables need to be represented as stochastic processes defined on some probability space and indexed by a parameter *t* ∈ [0, *T*] that represents the time. A natural formalism for describing the time evolution of such systems consists of the Itō stochastic differential equations (SDE) (8). For a closed system with two types of molecules *x* and *y*, the variables are the numbers of molecules denoted by *X(t)* and *Y(t)*, and the system of SDEs reads as:

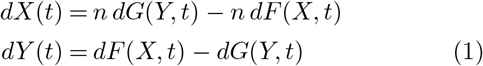

where *n* is the number of molecules of type *x* that make up *y.* The interconversion terms *dF* and *dG* are each expressed as the sum of a deterministic term with drift coefficient *F* and *G*, respectively, and of a stochastic term with diffusion coefficient 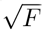 and 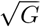:

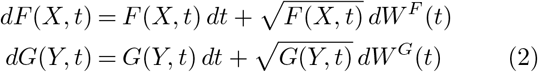

*W^F^(t)* and *W^G^(t)* stand for two independent Wiener processes: *W^F^(t)* − *W^F^(t’)* and *W^G^(t)* − *W^G^(t’)* follow a *N*(0,*t* − *t*’) distribution for all (*t,t*’), and *W^F^*(0) = 0 = *W^G^*(0). Wiener processes have continuous-valued realizations and are thus appropriate when the number of molecules is large enough to be approximated as a continuous variable. The Ito SDE formalism is equivalent, under some mild conditions, to the Fokker-Plank and master equation formalisms (8; 9).

For the simplicity of the subsequent calculations, we approximated the continuous SDEs given by Eqs (1,2) by discrete-time SDEs, where the time interval [0, *T*] is subdivided in Ξ equal-length intervals 0 = *t*_0_ < … < *t*_Ξ_ = *T*, with *t*_τ_ = τΔ*t* and Δ*t* = T/Ξ. Using the Euler-Maruyama discretization scheme (10), the discrete-time SDEs are:

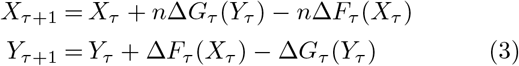

for all positive integers *τ* ∈ [0, Ξ]. The discretized interconversion terms are given by:

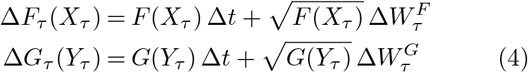

with *W_τ_* = *W t_τ_)* and Δ*W_τ_* = *W*_τ+1_ − *W*_τ_, so that in particular *W*_0_ = 0, *E*(Δ*W*_τ_) = 0 and **Var**(Δ*W*_τ_) = Δ*t*.

All systems considered here converge towards an equilibrium steady state in the long-time limit, obtained by first taking the limit *T* = ΞΔ*t* → co followed by Δ*t* → 0.

## 3. HOMO-OLIGOMERS

Consider the closed system depicted in Fig. 1a, in which *n* molecules of type *x* interact to form oligomers of type *y* without intermediate states. The discrete-time SDEs describing this system are given by Eqs (3-4) with:

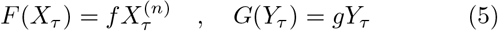

where 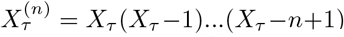. Since the system is closed, the combination of the two equations (3) yields the conservation constraint on the total number of molecules at all times:

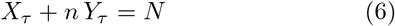

**Fig. 1.**
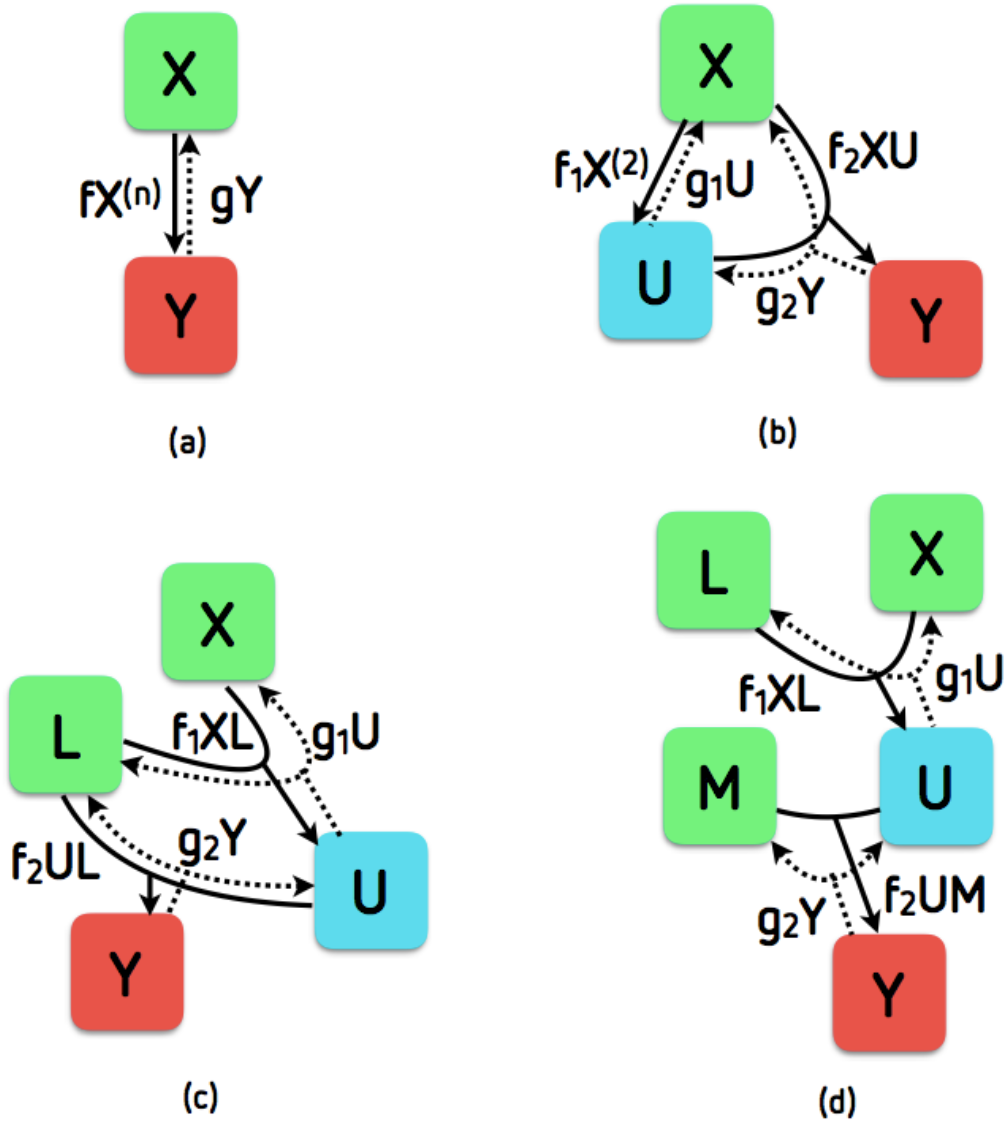
Schematical representation of the oligomerization systems studied in this paper. The green boxes correspond to monomers (*X, L, M*), the red boxes to oligomeric products (*Y*), and the blue boxes to intermediate states (*U*).

The mean of this equation and the mean of this equation multiplied by *X_τ_* or *Y_τ_* yield the relations:

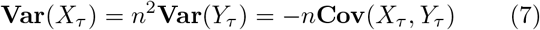

The covariance is thus always negative and equal to its lowest bound: 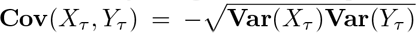. As a consequence, the conservation of the number of particles (6) implies the vanishing of the determinant of the covariance matrix: *C*(*X_τ_*, *Y_τ_*) = 0.

Another result following from Eq. (7) is that **Var**(*Y_τ_*) = **Var**(*X_τ_*) when *n* =1, and **Var**(*Y_τ_*) < **Var**(*X_τ_*) when *n* > 1; the variance ratio is equal to 1/*n*^2^. Thus, the higher the oligomer degree, the smaller the oligomer variance compared to the monomer variance.

To completely solve the model, we need to go back to Eqs (3-5), and take their mean and the mean of their squares at the steady state. This yields the additional relations:

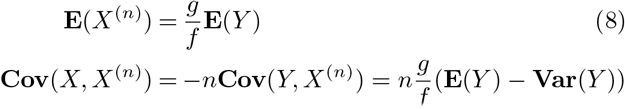

where the numbers of molecules at the steady state are denoted by *X* and *Y* (without subscript). Note that only the ratio of the parameters, *f*/*g*, enters in the equations.

For *n* = 1, the system of equations closes and we get:

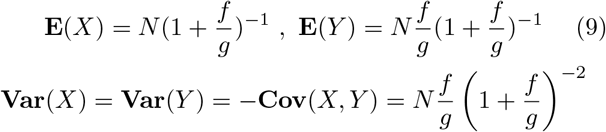

For *n* ≥ 2, the system does not close, and we had to make approximations to solve it. We used the moment closure approximation (12), assuming **E**(*X*) ≫ 1 and **E**(*Y*) ≫ 1. This yields:

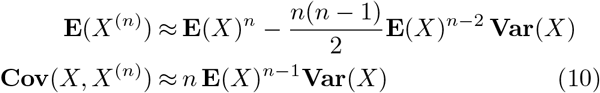

Eqs (8) now reduces to the closed system of algebraic equations:

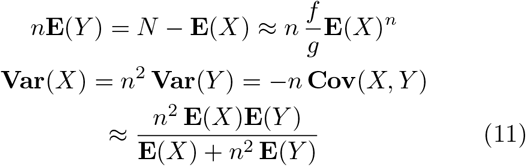

With these approximations, the mean, variance and co-variance of *X* and *Y* can be expressed as a function of the parameters. In a first step, we obtained a general expression of these quantities in terms of the parameter ratio *f*/*g* and **E**(*X*) for all values of *n*:

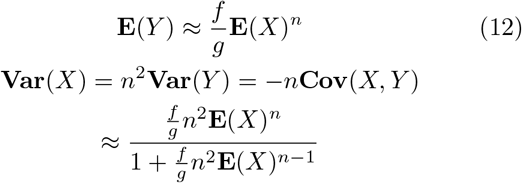

In a second step, **E**(*X*) can be written in terms of the parameters *f*/*g* and *N* in a *n*-dependent fashion. In particular, for *n* = 1 we recover Eqs (9), while for *n* = 2 we have:

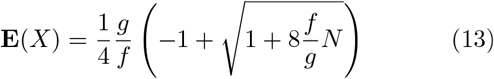

and for *n* =3:

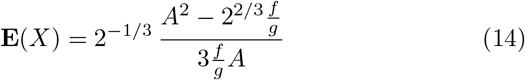

with 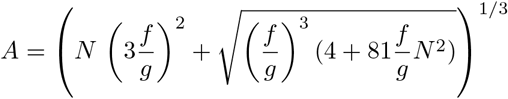

Thus, when *f*/*g* increases from 0 to ∞, **E**(*Y*) increases from 0 to *N* and **E**(*X*) decreases from *N* to 0, for all values of *n*. The variances **Var**(*X*), **Var**(*Y*) and the covariance –**Cov**(*X, Y*) vanish for *f*/*g* = 0 and *f*/*g* = ∞, and reach a maximum at 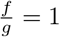 for *n* = 1, at

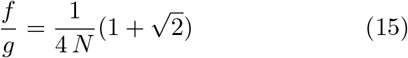

for *n* = 2 and at

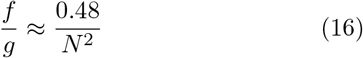

for *n* = 3. The mean and variances of *X* and *Y* as a function of *f*/*g* are shown in Fig 2, for fixed values of *N*. Clearly, they vary with the oligomer degree *n*.

**Fig. 2.**
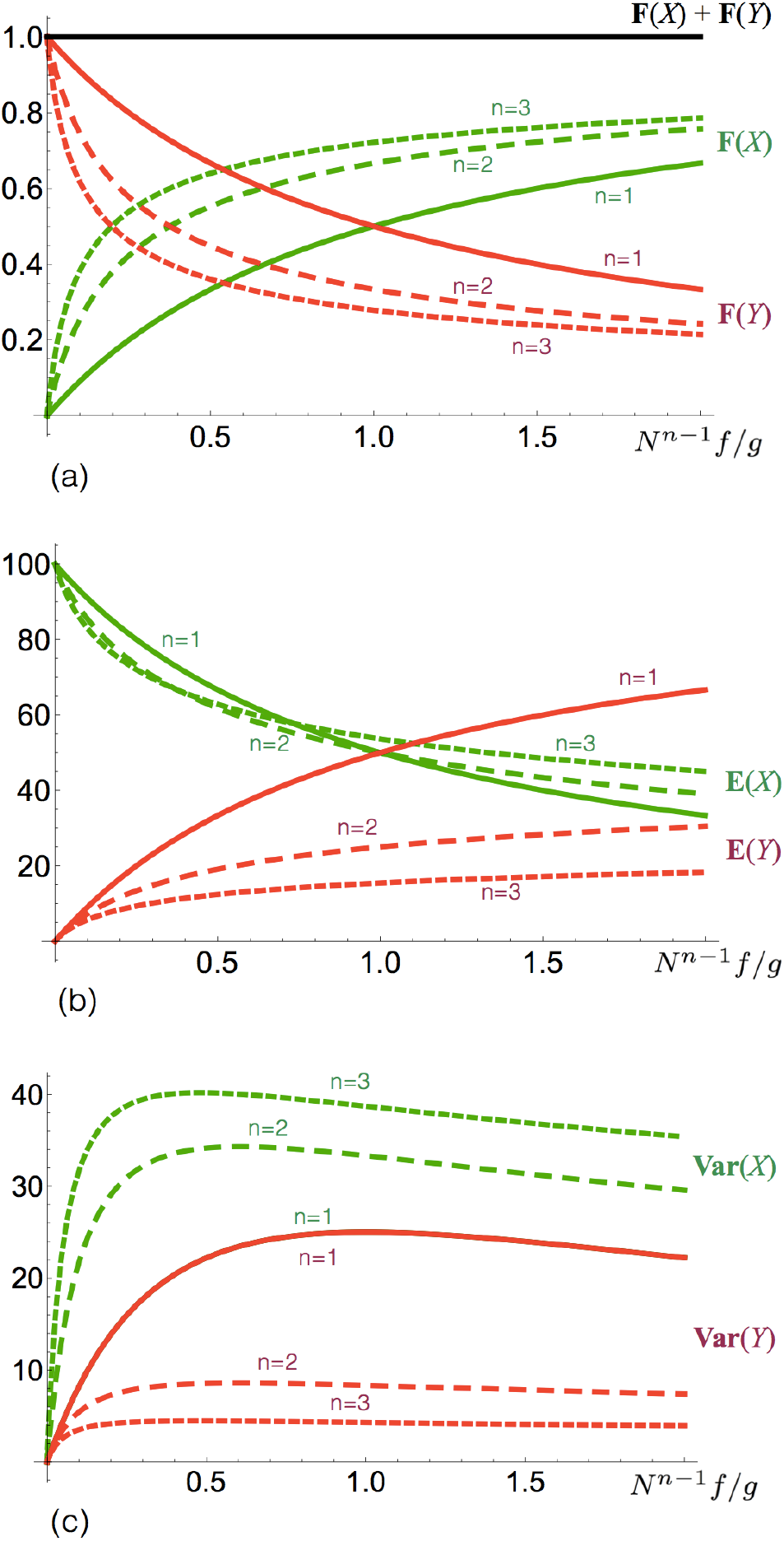
**Stochastic behavior of homo-oligomeration processes, as a function of the interconversion parameter** 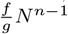, for oligomers of degree *n* = 1 to 3. N is taken equal to 100. (a) Fano factors; (b) Mean values; (c) Variances.

The intrinsic noise on a stochastic variable ***X*** is usually estimated by its Fano factor **F**, defined as the variance to mean ratio: **F(*X*) = Var(*X*)/E(*X*)**. This factor measures the deviation from Poissonian noise: when it is lower than one, the intrinsic noise is reduced compared to the Poisson distribution, and when it is larger than one, the noise is amplified.

From Eqs (13) follows that the sum of the Fano factors of the two molecular species x and y at the steady state is equal to one - the system’s rank - for all n, independently of the values of the parameters:

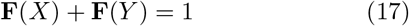

This result has been shown to hold for general deficiency zero systems (7)^1^. The Fano factors **F**(*X*) and **F**(*Y*) are thus lower than one and both molecular species exhibit sub-Poissonian noise. Strikingly, the *N*^*n*−1^*f*/*g* interval where the intrinsic noise is higher on the oligomer than on the monomer (**F**(*X*) < **F**(*Y*)) is reduced for larger oligomer degree *n* (see Fig. 2). Indeed the crossing point between the **F**(*X*) and **F**(*Y*) curves is 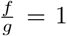 for *n* =1, 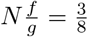 for *n* = 2 and 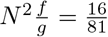 for *n* = 3.

Since the system is closed, we can define the reaction free energy as:

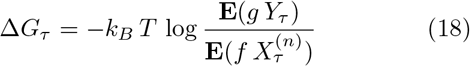

where *k_B_* is the Boltzmann constant and *T* the absolute temperature. This quantity vanishes at the steady state, which is in this case an equilibrium state. The standard free energy, Δ*G*^0^, is then:

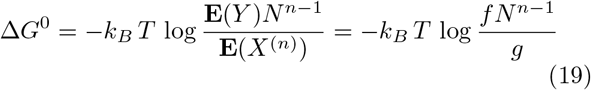

Thus, the larger the *f*/*g* ratio for fixed *N*, the more negative the standard free energy, and the stabler the product *y*.

## 4. HOMO-TRIMERS WITH DIMERIC INTERMEDIATE STATE

Consider now the case in which molecules *x* first form homodimers *u* and then homotrimers *y*, as depicted in Fig. 1b. This system corresponds to the homo-trimerization (*n* = 3) described in the previous section, but with an intermediate state. The discrete-time SDEs describing this system are:

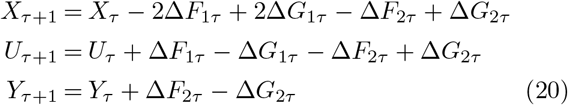

where

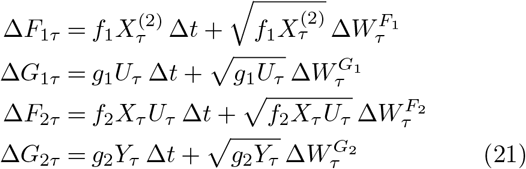

Δ*W*^*F*_l_^, Δ*W*^*F*_2_^, Δ*W*^*G*_l_^ and Δ*W*^*G*_2_^ are independent Wiener processes. The combination of these three equations, which imposes the conservation of the total number of particles at all times, is here:

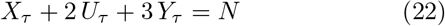

This conservation equation implies the following relations on the mean, variances and covariances:

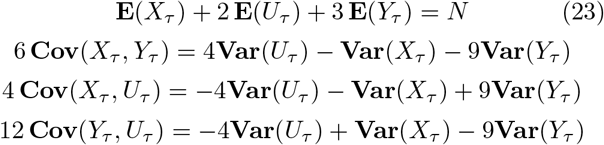

as well as the vanishing of the determinant of the covariance matrix: *C*(*X_τ_*, *Y_τ_*, *U_τ_*) = 0.

To fully solve equations Eqs (20,21) at the steady state, we used again the moment closure approximation (12). We got in this way all variances, covariances and mean values as a function of the parameter ratios *f*_1_ /*g*_1_ and *f*_2_/*g*_2_ and **E**(*X*):

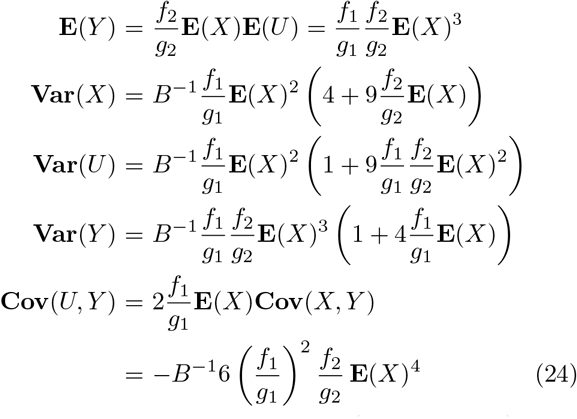

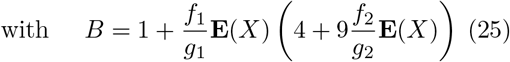

Here also, all the covariances are negative. These equations directly yield the Fano factor relation:

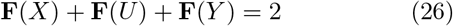

which generalizes Eq. (17) for systems of rank two (7).

To get **E**(*X*) in terms of the parameters, we had to solve the additional equation:

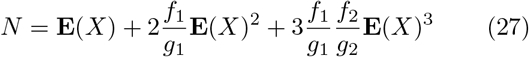

This yields:

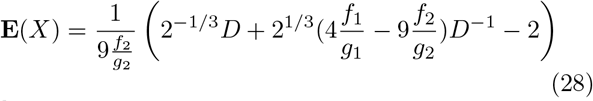

with

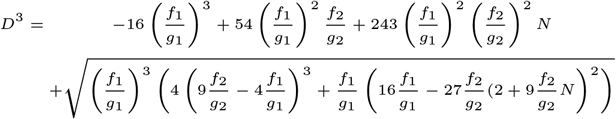

As illustrated in Fig. 3, the mean number of monomers and dimers **E**(*X*) and **E**(*U*) decrease with increasing parameter values *f*_2_/*g*_2_, for fixed *N* and *f*_1_/*g*_1_, whereas he number of trimers increase. In contrast, the variances show a maximum for some ranges of parameters. The Fano actors of the monomeric and dimeric states increase with *f*_2_/*g*_2_, whereas the Fano factor of the trimer decreases.

**Fig. 3.**
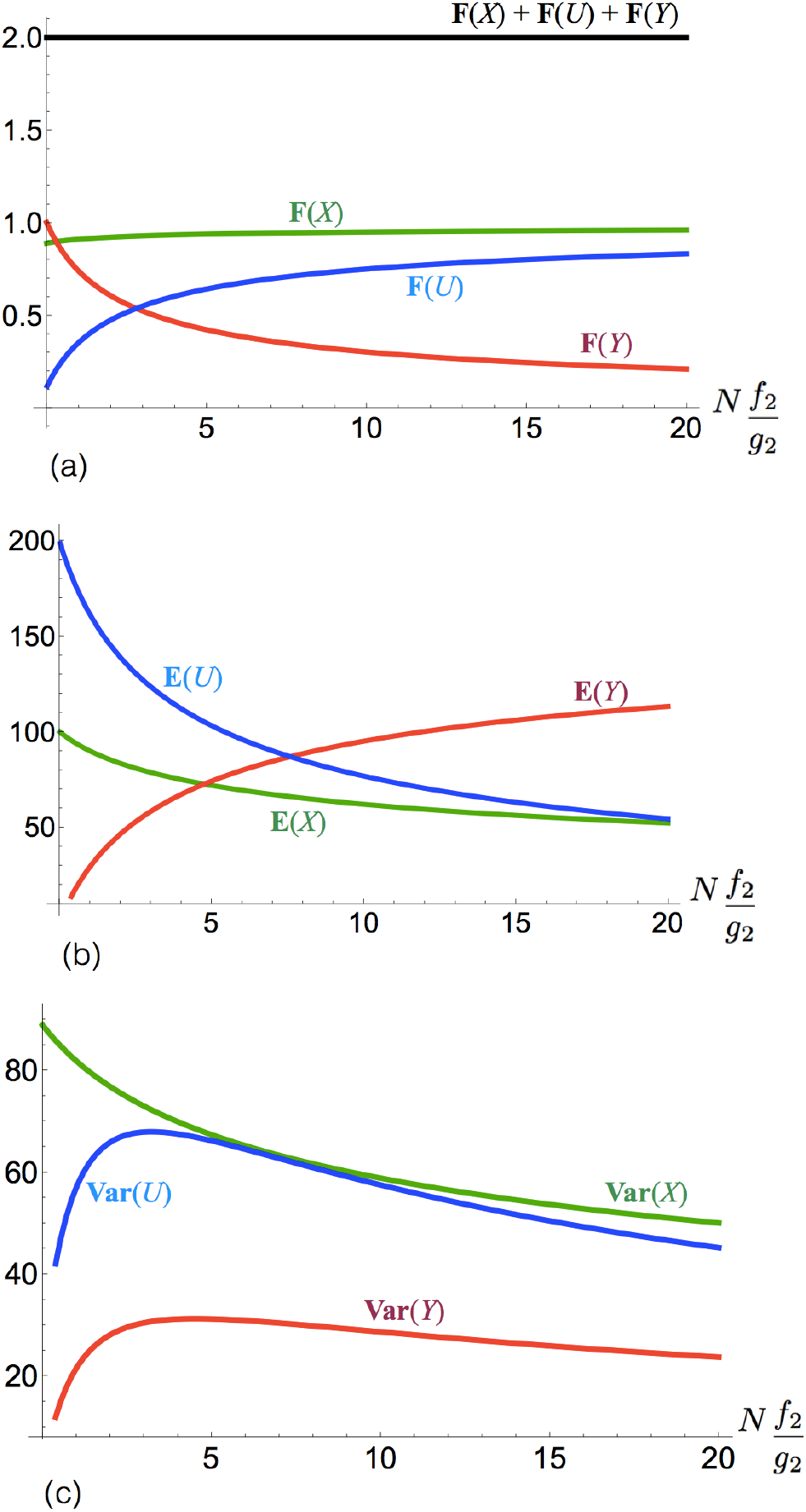
**Stochastic behavior of a homotrimerization process with a dimeric intermediate state**, as a function of *N f*_2_/*g*_2_. *N* is equal to 500 and *N f*_1_/*g*_1_ = 10. (a) Fano factors; (b) Mean values; (c) Variances.

The standard free energies of the two reactions are:

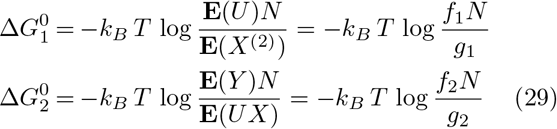

at the steady state. Cooperative and anticooperative processes consist of two or more binding reactions that are not independent, in which the formation of a first complex makes more or less likely its interaction with other molecules. Such phenomena are observed in a large variety of biochemical processes (13). In terms of free energy, the system described in this section displays a cooperative behavior when 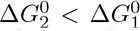, and an anticooperative behavior when 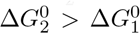. In terms of the cooperativity index defined as:

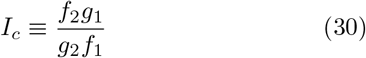

the system is cooperative when *I_c_* > 1 and anticooperative when *I_c_* < 1.

As shown in Fig. 4, the higher the cooperativity, the lower **F**(*Y*), and thus the lower the intrinsic noise on the reaction product. Moreover, for constant cooperativity index, the noise is reduced when increasing 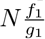.

**Fig. 4.**
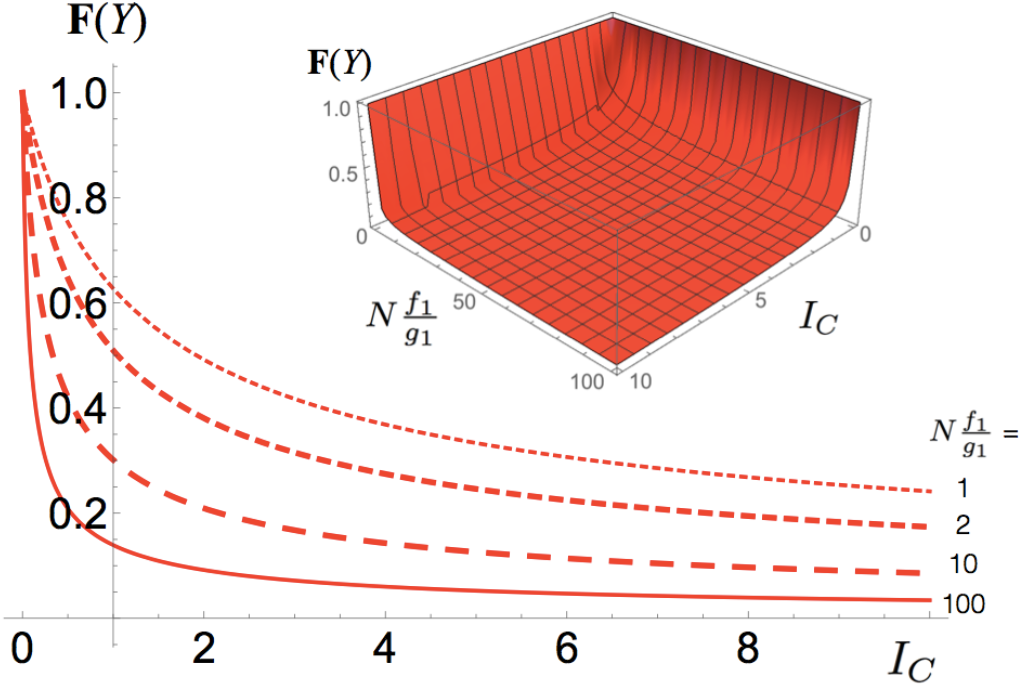
**Fano factor of the trimer **F**(*Y*) as a function of the cooperativity index** *I_C_* for different values of 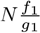 (2D graph), and as a function of *I_C_* and 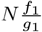 (3D graph); *N* is taken equal to 500.

## 5. (2,1)-HETERO-TRIMERS WITH DIMERIC INTERMEDIATE STATE

We generalized the trimerization of identical monomers described in the previous section to the case where one monomer *x* binds to another monomer ℓ to form an dimer *u*, which in turn binds to a second monomer ℓ to form a (2,1)-heterotrimer. This system is illustrated in Fig. 1c, and describes, for example, a protein *x* that binds successively to two ligands ℓ, leading to the protein-ligand complex *y*. The following discrete-time SDEs describe this system:

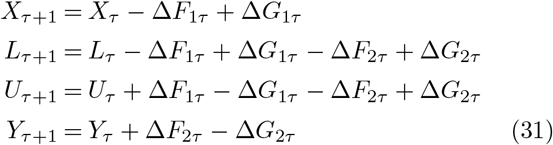

where

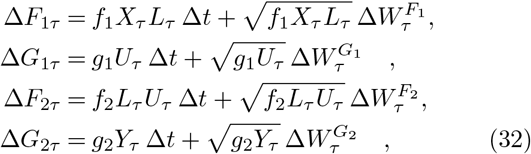

where Δ*W*^*F*_l_^, Δ*W*^*F*_2_^, Δ*W*^*G*_l_^ and Δ*W*^*G*_2_^ are independent Wiener processes.

Given that the system is closed, we get two relations, obtained as linear combinations of the four equations (31), which ensure the conservation of the number of molecules of species *x* and ℓ at all times. These are:

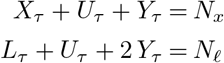

Using the same techniques as in the previous sections, we analytically solved the system of Eqs (31) under the moment closure approximation. We got in particular that the sum of the Fano factors of all molecular species is, as expected, equal to the system’s rank:

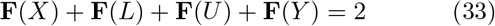

As illustrated in Fig. 5, the Fano factors of the monomer ℓ and of the intermediate state u increase with the parameter *f*_2_/*g*_2_, for fixed *f*_1_/*g*_1_, whereas the Fano factors of the monomer *x* and the trimer *y* decreases. Note that for other parameter values, **F**(*X*) also increases, whereas **F**(*Y*) always decreases. This behavior can again be related to the standard reaction free energies, which are in this case:

**Fig. 5.**
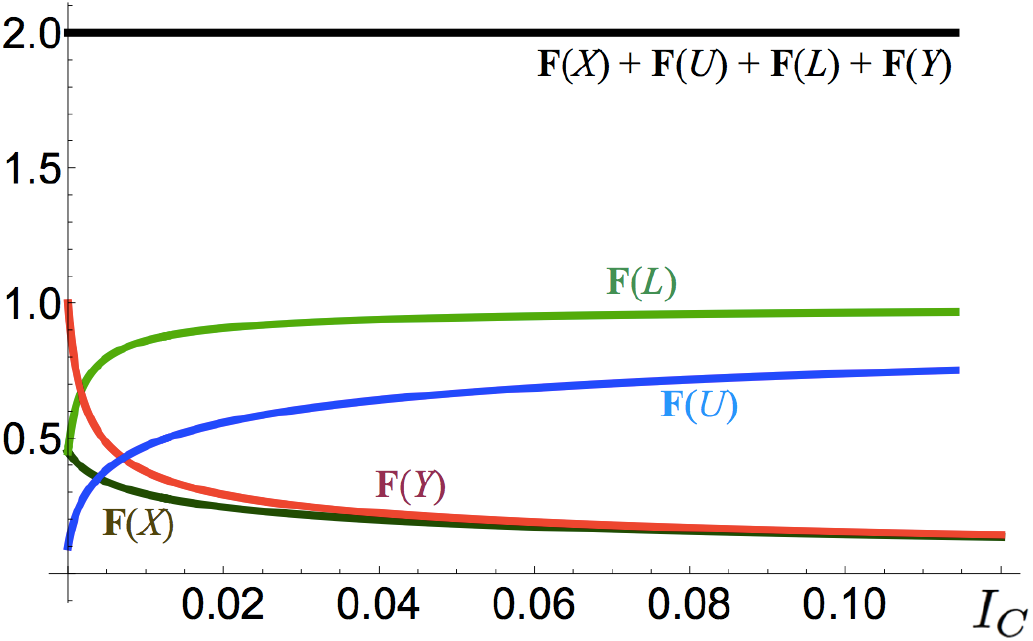
**Fano factors of (2,1)-heterotrimers, as a function of the cooperativity** *I_C_* with 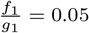 and *N_x_* = 500 = *N_ℓ_*.

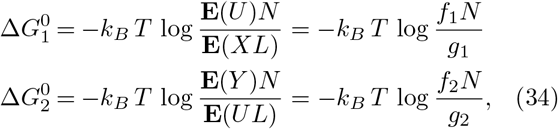

with *N* = *N_x_* + *N_ℓ_*. We found here also that the higher the cooperativity value *I_C_*, the lower the Fano factor on the trimers *y*.

## 6. (3)-HETEROTRIMERS WITH DIMERIC INTERMEDIATE STATE

We finally considered the trimerization of three different monomers *x*, ℓ and *m*, with an intermediate state u consisting of dimers built from *x* and £, as illustrated in Fig. 1d. Generalizing the equations and the approach used in the previous sections, we find that the molecular conservation relations imply the vanishing of the determinant of the covariance matrix at all times: *C*(*X_τ_*, *L_τ_*, *M_τ_*, *U_τ_*, *Y_τ_*) = 0, and the equality between the sum of the Fano factors of all molecular species and the rank of the system, independently of the parameter values:

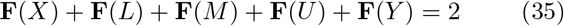

Furthermore, the Fano factor of the trimer y decreases with the increase of the cooperativity index *I_C_*, for fixed *f*_1_/*g*_1_. Hence, again, the higher the cooperativity, the larger the noise reduction on the complex.

## 7. CONCLUSION

On the basis of the closed homo-and hetero-oligomerization systems studied, we conclude that each molecular species is subject to sub-Poissonian noise, while the sum of their Fano factors is equal to the rank of the system. The intrinsic noise on the oligomeric product is all the more reduced as the oligomeric degree is high, and the reactions are cooperative.

There are still a several points that need to be addressed. The SDE formalism and moment closure approximation are only valid for sufficiently large numbers of molecules, and the validity of our results for small *N* need to be checked. To apply our approach to biological systems of interest, which are in general open systems interacting with the environment, our analysis need to be extended to such systems; our first results in this context are presented in (7; 14). A further stage will be the comparison of our analytical results with experimental data on fluctuations in targeted biochemical networks.

## Footnotes

⋆ FP and MR are research assistant and research director, respectively, at the Belgian Fund for scientific research (FNRS).

## Footnotes

1 The deficiency is a structural characteristic of the network defined as the number of complexes in the network minus the rank and number of connected components (11).

